# *Drosophila melanogaster* Toll-9 elicits antiviral immunity against *Drosophila* C virus

**DOI:** 10.1101/2024.06.19.599730

**Authors:** Manish Chauhan, Peter E. Martinak, Benjamin M. Hollenberg, Alan G. Goodman

## Abstract

The Toll pathway plays a pivotal role in innate immune responses against pathogens. The evolutionary conserved pathogen recognition receptors (PRRs), including Toll like receptors (TLRs), play a crucial role in recognition of pathogen associated molecular patterns (PAMPs). The *Drosophila* genome encodes nine Toll receptors that are orthologous to mammalian TLRs. While mammalian TLRs directly recognize PAMPs, most *Drosophila* Tolls recognize the proteolytically cleaved ligand Spätzle to activate downstream signaling cascades. In this study, we demonstrated that Toll-9 is crucial for antiviral immunity against *Drosophila* C virus (DCV), a natural pathogen of *Drosophila*. A transposable element insertion in the *Toll-9* gene renders the flies more susceptible to DCV. The stable expression of Toll-9 in S2 cells confers resistance against DCV infection by upregulation of the RNAi pathway. Toll-9 promotes the dephosphorylation of AKT, resulting in the induction of antiviral RNAi genes to inhibit DCV replication. Toll-9 localizes to the endosome where it binds dsRNA, suggesting its role to detect viral dsRNA. Toll-9 also induces apoptosis during DCV infection, contributing to its antiviral role. Together, this work identifies the role of Toll-9 in antiviral immunity against DCV infection through its ability to bind dsRNA and induce AKT-mediated RNAi antiviral immunity.

**IMPORTANCE:** Insects rely on innate immunity and RNA interference (RNAi) to combat viral infections. Our study underscores the pivotal role of *Drosophila* Toll-9 in antiviral immunity, aligning with findings in *Bombyx mori*, where Toll-9 activation upregulates the RNAi component *Dicer2*. We demonstrate that *Drosophila* Toll-9 functions as a pattern recognition receptor (PRR) for double-stranded RNA (dsRNA) during *Drosophila C* virus (DCV) infection, akin to mammalian TLRs. Toll-9 activation leads to the upregulation of key RNAi components, *Dicer2* and *Argonaute2*, and dephosphorylation of AKT triggers apoptosis via induction of proapoptotic genes *Hid* and *Reaper*. This study also reveals that Toll-9 localizes in endosomal compartments where it interacts with dsRNA. These insights enhance our understanding of *Drosophila* innate immune mechanisms, reflecting the evolutionary conservation of immune responses across diverse species and providing impetus for further research into the conserved roles of TLRs across the animal kingdom.

## INTRODUCTION

The innate immune system is the first line of defense against invading pathogens, and since insects lack adaptive immunity, they solely rely on innate immunity (Agaisse 2007). The innate immune system is an efficient and fast acting defense mechanism that identifies pathogen associated molecular patterns (PAMPs) through pathogen recognition receptors (PRRs) (Lemaitre et al. 1996; Litman, Rast, and Fugmann 2010). Upon interacting with PAMPs, PRRs initiate downstream signaling to mount an antimicrobial response (Lin, Cohen, and Wasserman 2020; Lindsay and Wasserman 2014). In higher vertebrates such as mammals, Toll-like receptors (TLR) recognize the pathogen associated molecular patterns (PAMPs) such as lipopolysaccharide (LPS), Fungal zymosan, and cytosolic nucleic acids (Akira and Takeda 2004; Iwasaki and Medzhitov 2004; Kawai and Akira 2010).

The Toll receptor was first discovered as a type I transmembrane receptor for its role in dorsoventral patterning during embryonic development in *Drosophila*. Later, its role in innate immunity was deciphered (Anderson, Jürgens, and Nüsslein-Volhard 1985; Lemaitre et al. 1996). *Toll-1* was identified as an important immune receptor for its role in activating antimicrobial peptides (AMPs) against gram positive bacteria and fungi. There are nine *Toll* genes in *Drosophila*, and *Toll-1* has a major role in innate immune signaling (Valanne et al. 2022; Valanne, Wang, and Rämet 2011). Furthermore, studies suggest Toll-8 (Tollo) mediates immunity in the trachea of *Drosophila*, and Toll-7 regulates anti-viral responses independent of an NF-κB signaling pathway (Akhouayri et al. 2011; Nakamoto et al. 2012). However, the immunological function of the remaining *Drosophila* Toll receptors has not yet been deciphered (Anthoney, Foldi, and Hidalgo 2018; Tauszig et al. 2000).

Of the nine *Drosophila* Tolls, Toll-9 is most closely related to mammalian TLRs and is orthologous to TLR10 (Anthoney, Foldi, and Hidalgo 2018). Toll-9 lacks multiple autoinhibitory cysteine-rich motifs which differentiates it from other Toll receptors and makes it more similar to mammalian TLRs (Khush et al. 2000; Imler and Zheng 2004). Previous studies on Toll-9 suggest its role in innate immunity, and Ooi et al. has shown that overexpression of Toll-9 induces production of AMPs (Ooi et al. 2002). However, Narbonne-Reveau et al. demonstrated that in the absence of Toll-9 there was no change in basal induction levels of AMPs and Toll-9 overexpression was unable to induce AMPs production during oral bacterial infection (Narbonne-Reveau, Charroux, and Royet 2011). This finding was contrary to the study by Ooi et al., which showed that Toll-9 induces basal levels of AMPs and that its overexpression further increases AMP production. Recent studies have highlighted the role of Toll-9 in cell fitness. Specifically, the activation of Toll-9 signaling has been shown to induce apoptosis during cell competition in developing tissues, thereby eliminating unfit cells and promoting proliferation in surviving cells (Meyer et al. 2014; Shields et al. 2022). Furthermore, Toll-9 has been shown to regulate aging and neurodegenerative processes in a *Drosophila* model of tauopathy (Sakakibara et al. 2023).

Arthropods rely on the recognition of virus derived double-stranded (dsRNA) to activate the RNAi pathway and mount antiviral responses (Cherry and Silverman 2006; de Faria et al. 2013). *Drosophila* C virus (DCV) is a positive-sense RNA virus of the family Dicistroviridae and is one of the most well-described *Drosophila* viruses. Since *Drosophila’s* major antiviral pathway is mediated through RNAi, and DCV generates dsRNA replication intermediates that are detected by the RNAi components such as *Dicer2* and *Argonaut2* to cleave the viral dsRNA (Karlikow, Goic, and Saleh 2014; Bronkhorst et al. 2012; Kemp et al. 2013), we hypothesized that *Drosophila* encodes a PRR that binds to dsRNA during DCV infection. Since previous studies showed that TLR10 recognizes dsRNA in the endosome and compete for the dsRNA binding with TLR3 (S. M.-Y. Lee et al. 2018), and Toll-9 is orthologous to TLR10, we further hypothesized that Toll-9 is a PRR for dsRNA during DCV infection. As such, we found that a transposable element insertion in *Toll-9* renders flies more susceptible to DCV infection compared to isogenic control flies. We expressed Toll-9 in S2 cells and found that it confers resistance against DCV infection by upregulating *Dicer2* and *Argonaut2* expression levels. We observed that Toll-9 colocalizes with Rab5 (early endosome) and Rab7 (late endosome) in the endosome and binds to dsRNA. We also found Toll-9 induces apoptosis during DCV infection by inducing expression of proapoptotic gene *Hid* and *Reaper*. Altogether, our data suggest that Toll-9 plays a crucial role in antiviral immunity by inducing RNAi pathway components and apoptosis.

## RESULTS

### Toll-9 mutant flies display increased susceptibility to DCV infection

We investigated the role of Toll-9 in *Drosophila* antiviral defense, by infecting isogenic control *w^1118^* and transposable element insertion mutant *Toll-9^0024-G4^* flies with DCV. *Toll-9^0024-G4^* mutant flies are more susceptible to DCV infection because they exhibit higher mortality rates compared to *w^1118^* isogenic control flies (Fig. 1A). Next, we examined DCV replication in mutant *Toll-9^0024-G4^* flies compared to control *w^1118^* flies by measuring DCV RNA and capsid protein levels. Our analysis revealed that the levels of DCV *replicase ORF1* mRNA were significantly higher in *Toll-9^0024-G4^* flies at both 1- and 3-days post-infection (dpi) compared to the control *w^1118^*flies (Fig. 1B). We also observed elevated levels of the DCV capsid protein in *Toll-9^0024-G4^* flies infected with DCV at both 1- and 3 dpi, further indicating increased viral replication in the absence of functional Toll-9 (Fig. 1C). Additionally, we quantified the levels of infectious virus in DCV-infected control *w^1118^*and mutant *Toll-9^0024-G4^*. Our results indicate that DCV titers were higher in *Toll-9^0024-^*^G*4*^ than *w^1118^* flies at both 1- and 3-dpi (Fig. 1D). Together, the results suggest that *Toll-9* is involved in antiviral immunity against DCV infection in *Drosophila* and mutation in *Toll-9* renders flies more susceptible to DCV infection and exhibit increased viral load.

**Figure 1.**
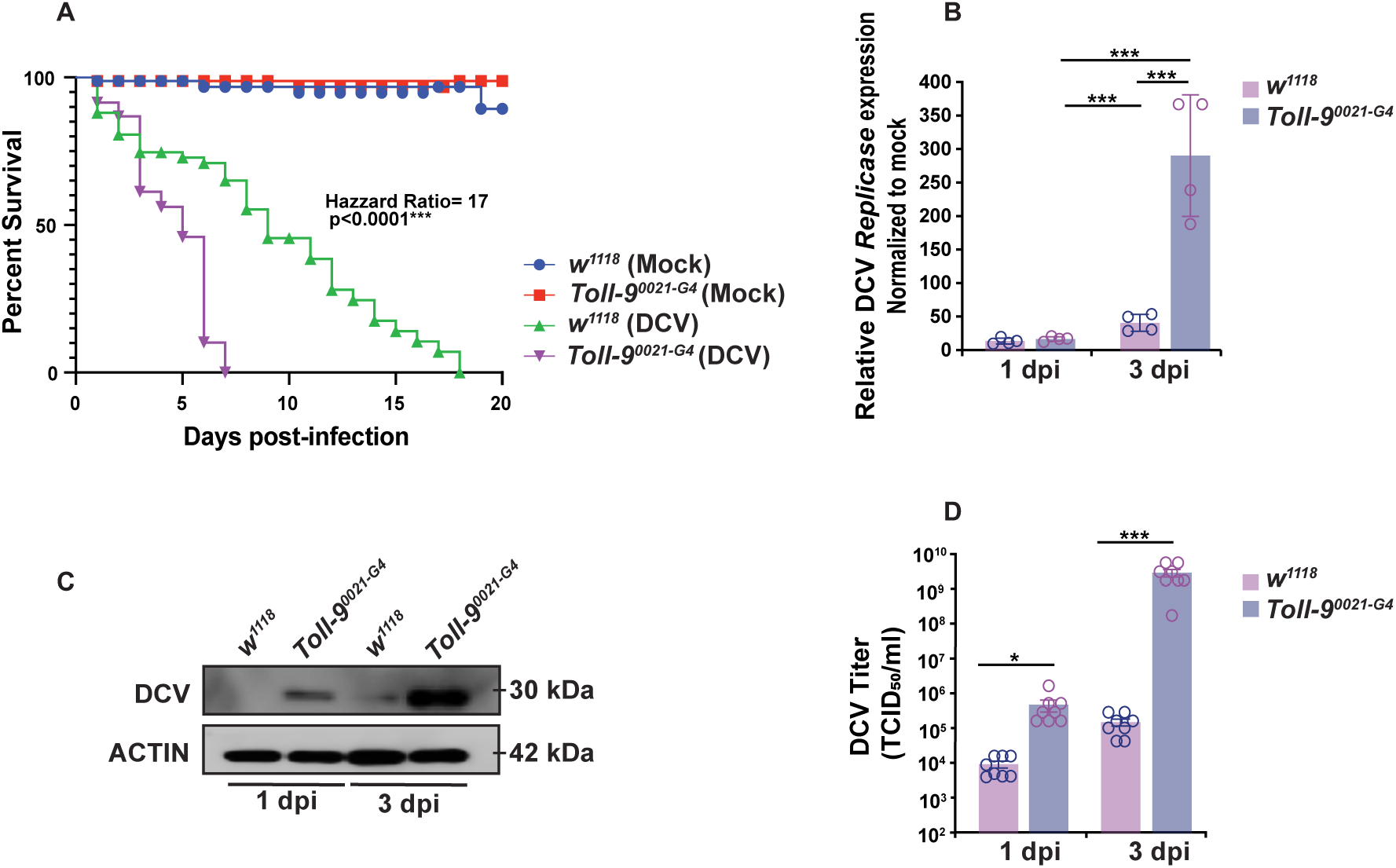
Toll-9 mutant flies display increased susceptibility to DCV infection. **(A)** Survival analysis for isogenic control *w^1118^* flies and *Toll-9^0024-G4^* flies injected with PBS and DCV. (N=80 flies from three independent experiments) **(B)** qPCR analysis DCV infected *w^1118^* flies and *Toll-9^0024-G4^* flies at 1 dpi and 3 dpi for DCV mRNA. Data are representative of four biological replicates (groups of five flies) each from three independent experiments. Error bars, SEM. Unpaired T test, *p<0.05; ***p<0.001. **(C)** Western blot analysis to probe DCV capsid protein with anti-DCV antibody from fly lysate of isogenic control *w^1118^* flies and *Toll-9^0024-G4^* flies injected with DCV at 1 dpi and 3 dpi. Data are representative of three independent experiments. **(D)** DCV titer in S2 cells infected with fly lysate from isogenic *w^1118^* flies and *Toll-9^0024-G4^* flies injected with DCV calculated as 50% Tissue culture infectious dose (TCID_50_/ml). Data are representative of eight biological replicates (groups of five flies) from four independent experiments. Error bars, SEM. Unpaired T test, *p<0.05; ***p<0.001.

### *Toll-9* regulates the expression of RNAi-dependent genes, *Dicer2* and *Argonaute2*

To address the role of Toll-9 in antiviral immunity and the control of DCV infection *in vivo*, we examined DCV-infected *w^1118^*isogenic control and transposable element insertion mutant *Toll-9^0024-G4^*flies by qRT-PCR. We observed that DCV infected mutant *Toll-9^0024-G4^*flies have lower expression of RNAi pathway genes *Dcr2* and *Ago2* (Fig. 2A-B). The JAK/STAT pathway also provides antiviral defense in *Drosophila* (Tafesh-Edwards and Eleftherianos 2020). During viral infection, the JAK/STAT pathway is activated when the upd2 and upd3 ligands bind to the Domeless receptor to activate STAT92E and induce expression of *Vir1* (Myllymäki and Rämet 2014; Ahlers et al. 2019). During DCV infection, *Toll-9^0024-G4^*mutant flies exhibited an elevated expression of JAK/STAT pathway-related genes compared to *w^1118^* control flies (Fig. 2C-E). It was previously shown that while the JAK/STAT pathway is an indicator of DCV infection, it was not sufficient for triggering an antiviral response to DCV (Dostert et al. 2005). Similarly, our results show that the increase of JAK/STAT signaling in DCV-infected *Toll-9^0024-G4^* mutant flies cannot compensate for the loss of RNAi in these mutant flies, which is the driver for increased DCV replication and susceptibility to infection.

**Figure 2.**
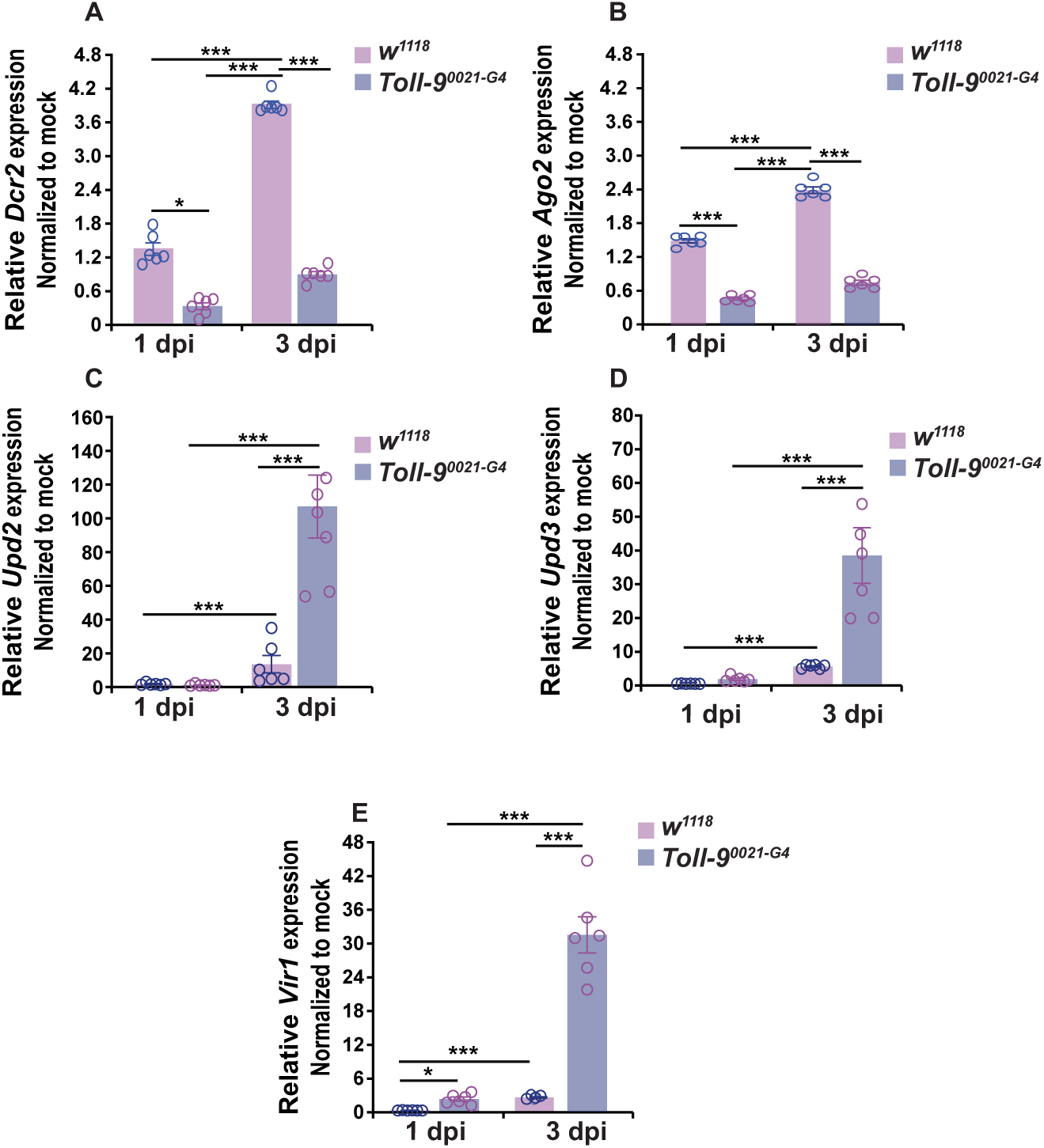
*Toll-9* regulates the expression of RNAi-dependent genes, *Dicer2* and *Argonaute2*. **(A-B)** qPCR analysis DCV infected *w^1118^* flies and *Toll-9^0024-G4^* mutant flies at 1 dpi and 3 dpi for RNAi pathway (*Dicer2* and *Arganoute2*). **(C-E)** qPCR analysis DCV infected *w^1118^* flies and *Toll-9^0024-G4^* flies at 1 dpi and 3 dpi for JAK/STAT pathway (*Upd2*, *Upd3* and *Vir1*). Data are representative of six biological replicates (groups of five flies) from three independent experiments. Error bars, SEM. Unpaired T test, *p<0.05; ***p<0.001.

### Stable expression of *Toll-9* reduces DCV infection in *Drosophila* S2 cells

To further elucidate the role of Toll-9 in defense against *Drosophila* C virus (DCV) infection, we generated a stable S2 cell line expressing Toll-9 under the control of a copper inducible promoter (Fig. 3A). The generation of a cell line stably expressing Toll-9 was necessary since modENCODE *Drosophila* cell line expression data deposited at FlyBase shows that *Toll-9* expression levels in S2 cells are extremely low to none (Gerstein et al. 2010; Brown et al. 2014; Cherbas et al. 2011; Gerstein et al. 2010). Upon DCV infection, Toll-9-expressing S2 cells displayed enhanced resistance compared to naïve S2 cells at 1 dpi, when induced with copper sulfate (CuSO_4_) (Fig. 3B). To investigate whether Toll-9’s antiviral effect is mediated through the activation of the RNA interference (RNAi) pathway, we assessed the expression of the RNAi pathway gene, *Dcr2*, in DCV-infected cells. CuSO_4_-induced Toll-9-expressing S2 cells exhibited a significantly elevated expression of *Dcr2* compared to naïve S2 cells, indicating Toll-9’s involvement in activating the RNAi pathway (Fig. 3C). Overall, our findings suggest that Toll-9 plays a crucial role in antiviral defense during DCV infection of S2 cells by restricting viral replication, through the activation of RNAi.

**Figure 3.**
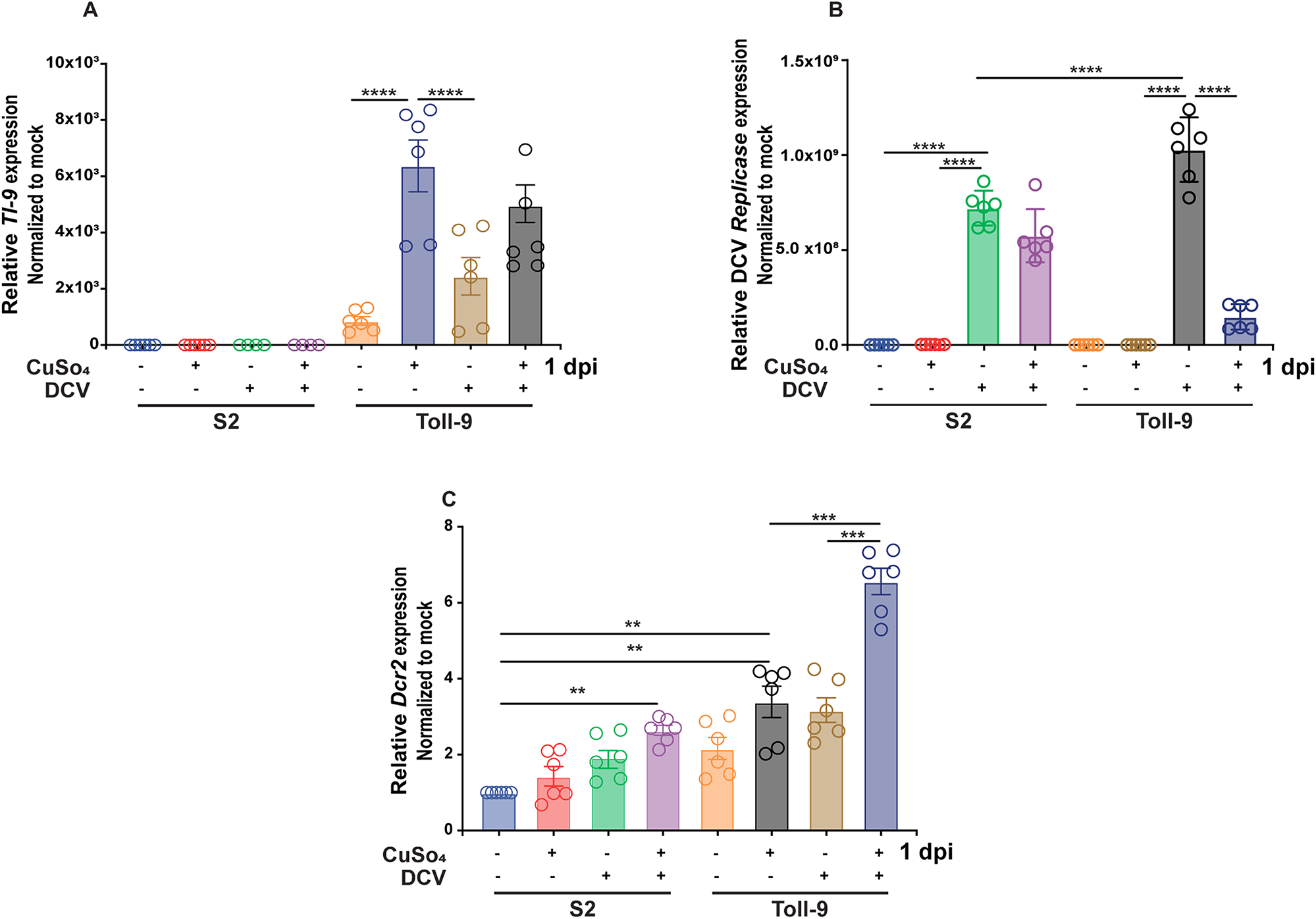
Stable expression of *Toll-9* reduces DCV infection in *Drosophila* S2 cells. **(A)** qPCR for detection of expression of *Toll-9* transcripts in mock- and DCV-infected S2 naive and S2 cells stably expressing TOll-9 in presence and absence of CuSO_4_ at 1 dpi. **(B-C)** qPCR analysis of DCV infected S2 naive and S2 cells expressing TOll-9 in presence and absence of CuSO_4_ at 1 dpi to detect expression of DCV *replicase ORF 1* transcript and *Dicer2* mRNA. Data are representative of six biological replicates (well of cells) from three independent experiments. Error bars, SEM. Unpaired T test, **p<0.01; ***p<0.001; ****p<0.0001.

### Toll-9 controls DCV infection via dephosphorylation of AKT

AKT (Protein Kinase B) is serine/threonine-specific protein kinase that plays crucial role in multiple cellular processes: glucose metabolism, cell cycle progression and protein synthesis (Hers, Vincent, and Tavaré 2011). During viral infections, viruses hijack various cellular pathways to facilitate their own survival and replication. Numerous studies on different types of viruses, including DNA viruses, RNA viruses, and retroviruses, have demonstrated that these pathogens manipulate the PI3K/AKT pathway. This manipulation leads to the regulation of apoptosis, alteration of splicing mechanisms, modulation of endocytosis, enhancement of RNA synthesis, and reorganization of the actin cytoskeleton (Cooray 2004; Ji and Liu 2008; Dunn and Connor 2012; Diehl and Schaal 2013; Swevers, Liu, and Smagghe 2018; Blanco et al. 2020). Previous studies on Sindbis virus (SINV) replication in *Drosophila* suggest its dependence upon the levels of AKT expressed in the cells underscores the significance of AKT in the viral life cycle. Intriguingly, SINV infection has been shown to induce AKT phosphorylation. This increase in AKT phosphorylation may contribute to the facilitation of SINV replication by promoting cellular processes conducive to viral proliferation (Patel and Hardy 2012). A recent study on West Nile Virus (WNV) infection in *Drosophila* S2 cells indicates a significant role for AKT phosphorylation during the infection process. The findings suggest that insulin activated phosphorylation of AKT during WNV infection leads to the inhibition of RNA interference (RNAi) and the activation of the JAK/STAT pathway to compensate the loss of RNAi induction to restrict WNV replication (Ahlers et al. 2019). In *Drosophila* S2 cells, the exact role of AKT phosphorylation during DCV infection has not been explored. Since AKT plays an important role in immune response and inflammation, we hypothesized that AKT could be involved in host cellular responses to DCV infection. We observed increased AKT phosphorylation during DCV infection of naïve S2 cells whereas the phosphorylation of AKT was reduced in the presence of Toll-9 (Fig. 4A). To further validate that AKT phosphorylation regulates DCV infection, naïve S2 and Toll-9 expressing S2 cells were treated with AKT VIII inhibitor for 16 hours followed by DCV infection. Pharmacological inhibition of AKT phosphorylation drastically reduced viral load from the DCV infected S2 and Toll-9 expressing S2 cells (Fig. 4B). We also examined cells using fluorescence microscopy to show the effect of Toll-9 on intracellular viral load in both the presence and absence of AKT inhibitor. Our results suggest that in the absence of AKT inhibitor, CuSO_4_-induced Toll-9 expressing S2 cells had lowest viral load compared to uninduced Toll-9 expressing S2 cells or naïve S2 cells (Fig. 4C). In the presence of AKT inhibitor, intracellular viral load was reduced in both cell types and treatment conditions (Fig. 4D). Next, to observe levels of infectious virus, cell culture supernatants were collected from DCV-infected naïve S2 cells and Toll-9 expressing S2 cells, both in the presence and absence of an AKT inhibitor. Our results show that the induction of Toll-9 expression or AKT inhibition resulted in suppression of DCV replication (Fig. 4E). Taken together, these results suggest Toll-9 restricts DCV replication by reducing dephosphorylation of AKT during infection.

**Figure 4.**
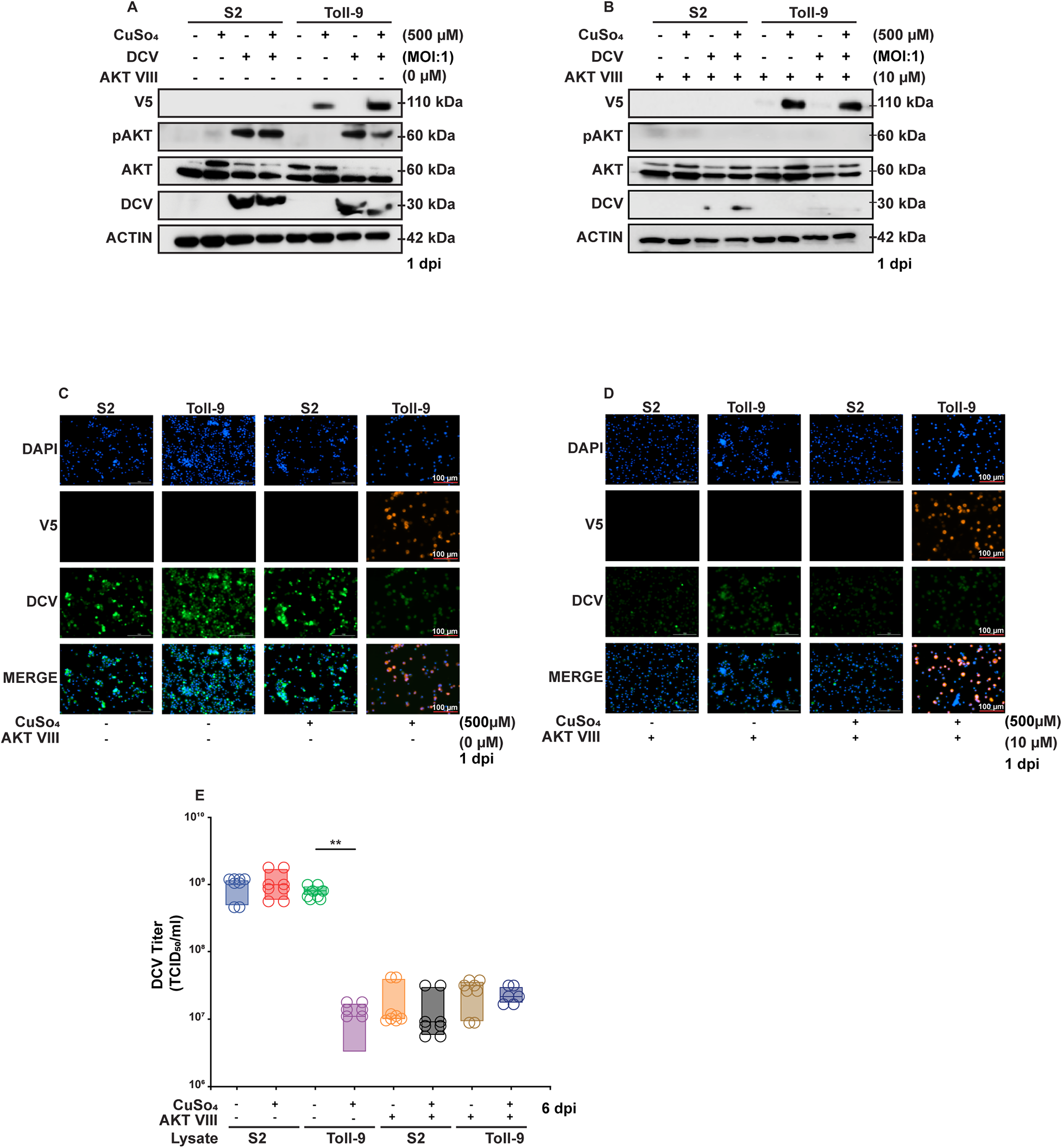
Toll-9 controls DCV infection via dephosphorylation of AKT. **(A-B)** Western blot analysis for indicated antibodies of cell lysate from DCV infected S2 cells and Toll-9 OE cells in presence and absence of CuSO_4_ at 1 dpi in presence and absence of Akt inhibitor. Data are representative of three independent experiments. **(B-C)** Micrograph shows presence of DCV in S2 naive cells and S2 cells expressing Toll-9 in presence and absence of CuSO_4_ at 1 dpi in presence and absence of Akt inhibitor. Data are representative of three independent experiments. **(E)** Viral titer of DCV infected S2 and Toll-9 OE cells culture supernatant in presence and absence of Akt inhibitor calculated as 50% Tissue culture infectious dose (TCID_50_/ml). Data are representative of eight biological replicates (well of cells) from four independent experiments. Error bars, SEM. Unpaired T test, **p<0.01.

### Subcellular localization of Toll-9 and interaction with dsRNA

To investigate the interaction of Toll-9 with dsRNA, we first explored the subcellular localization of Toll-9 in S2 cells. Toll-9 is a transmembrane protein, suggesting it may localize to intracellular membranes. We performed *in silico* analysis using the SignalP 6.0 database (Teufel et al. 2022) and identified the presence of a signal peptide. Specifically, Toll-9 contains a signal peptide belonging to Sec/SpI pathway that targets vesicular pathways with high probability score of 0.9. The predicted signal peptide is located in the N-terminal region of Toll-9 and exhibits characteristic features such as an N-terminal sequence, a hydrophobic core, a C-terminal region, and a cleavage site for Signal Peptidase I (SPI) (Fig. 5A).

**Figure 5.**
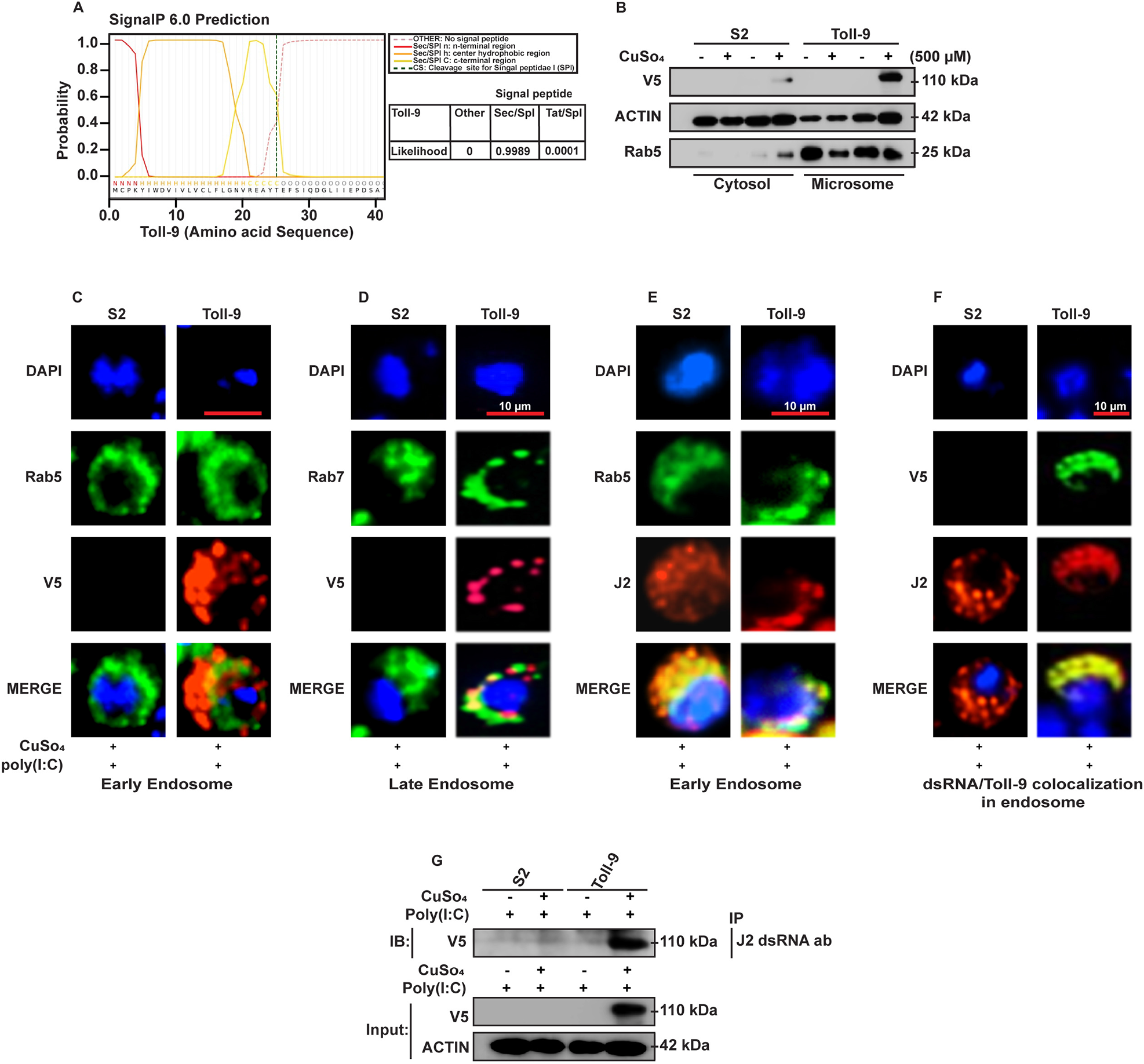
Subcellular localization of Toll-9 and interaction with dsRNA. **(A)** *In silico* prediction of signal peptide in Toll-9 protein sequence. Red solid line indicates predicted n-terminal region, orange solid line indicates the predicted center hydrophobic region, and yellow solid line indicates predicted c-terminal region of signal peptide. Black dotted line indicates the cleavage site (CS) of the signal peptide. Sec/SPI: Sec translocon transported secretory signal peptide/Signal Peptidase I Tat/SPI: Tat translocon transported Tat signal peptides/Signal Peptidase I **(B)** Western blot analysis demonstrating the presence of Toll-9/V5 in endosomes. Endosomal fractions were identified using Rab5 as a microsomal marker, while Actin served as a cytosolic marker. Data are representative of three independent experiments. **(C)** Micrographs show colocalization of Rab5-early endosome marker (green) and Toll-9 (anti-V5 tag ab-Red) in Poly(I:C) and CuSO_4_ (500 µM) treated Toll-9 OE and S2 cells. **(D)** Micrographs show colocalization of Rab7-Late endosome marker (green) and Toll-9 (anti-V5 tag ab-Red) in Poly(I:C) and CuSO_4_ (500 µM) treated Toll-9 OE and S2 cells. **(E)** Micrographs show colocalization of Rab5-early endosome marker (green) and Poly(I:C) (J2 anti-dsRNA ab-Red) in Poly(I:C) and CuSO_4_ (500 µM) treated Toll-9 OE and S2 cells. **(F)** Toll-9 (anti-V5 tag ab-Green) and Poly(I:C) (J2 anti-dsRNA ab-Red) in Poly(I:C) and CuSO_4_ (500 µM) treated Toll-9 OE and S2 cells. **(G)** Western blot analysis using the indicated antibodies following immunoprecipitation of V5 tag (Toll-9) using J2 dsRNA antibody from the lysate of Poly (I:C) treated Toll-9 OE and S2 cells in presence and absence of CuSO_4_ (500 µM). Data are representative from three independent experiments.

Next, we collected cellular fractions of naïve S2 and Toll-9-expressing S2 cells in presence and absence of CuSO_4_. We performed western blot analysis on cytosolic and microsomal fractions of induced S2 and Toll-9 expressing S2 cells to detect the presence of Toll-9 protein in cytosolic or microsomal fractions. Our results show that Toll-9 is predominantly present in microsomal fraction, suggesting its localization either in the ER (endoplasmic reticulum) or in vesicular system such as endosomes (Fig. 5B). To visualize the intracellular localization of Toll-9 we performed fluorescence microscopy using Rab5 (early endosomal) and Rab7 (late endosomal) markers to localize Toll-9. Both early and late endosomal markers Rab5 and Rab7 colocalize with Toll-9, suggesting its presence in the endosomal compartment (Fig. 5C-D). RNA viruses generally produce dsRNA as their replication intermediate during infection cycle irrespective of their genome type (DeWitte-Orr and Mossman 2010; Son, Liang, and Lipton 2015; Mondotte and Saleh 2018). As such, we treated S2 cells with the dsRNA analogue poly (I:C), and our results suggest that dsRNA was localized within the endosomes of both S2 cells and Toll-9 expressing S2 cells, as detected using the J2 dsRNA antibody and the endosomal marker Rab5 (Fig. 5E). Interestingly, we found that Toll-9 colocalized with dsRNA as Toll-9 and poly (I:C) were present in the endosome (Fig. 5F), suggesting that they might be interacting with each other.

To examine the interaction between dsRNA poly (I:C) and Toll-9, we performed immuno-pulldown assay using the J2 antibody with lysates from poly (I:C)-treated, Toll-9 expressing S2 cells in presence and absence of CuSO_4_. The J2 antibody effectively immunoprecipitated V5-tagged Toll-9 from the lysate of CuSO_4_-induced, poly (I:C)-treated, Toll-9-expressing S2 cells but not from naïve nor untreated S2 cells, suggesting that Toll-9 interacts with dsRNA (Fig. 5G). Moreover, since our results suggest that Toll-9 is localized in endosomes and interacts poly (I:C), Toll-9 may be activated by dsRNA during DCV infection of S2 cells.

### Toll-9 induces apoptosis during DCV infection of S2 cells

Cell death initiated during apoptosis is crucial for antiviral mechanism in multicellular organisms where it acts as a defense mechanism by preventing virus from further infecting the cells (Nainu et al. 2015; Nainu, Shiratsuchi, and Nakanishi 2017; Nainu et al. 2023; Roulston, Marcellus, and Branton 1999). Previous studies suggest that S2 cells undergo apoptosis upon DCV infection as an antiviral response, and apoptotic S2 cells are readily cleared by phagocytosis (Nainu et al. 2015). Our data suggests that Toll-9 reduces the viral load in S2 cells, so we next investigated whether Toll-9 confers an antiviral state in S2 cells by inducing apoptosis. Our results suggest that Toll-9 induces higher expression of proapoptotic genes *Hid* and *Reaper* during DCV infection at 1 dpi (Fig. 6A-B). To confirm that Toll-9 mediated induction of proapoptotic genes *Hid* and *Reaper* during DCV infection induces apoptosis, we performed an Annexin V assay to determine the apoptotic cell population in DCV infected or mock S2 cells, and Toll-9 expressing S2 cells in presence and absence of CuSO_4_ at 1 dpi. Our results revealed that Toll-9 expression increases apoptosis in S2 cells during DCV infection (Fig. 6C), suggesting that Toll-9 plays a role in inducing apoptosis during DCV infection in S2 cells.

**Figure 6.**
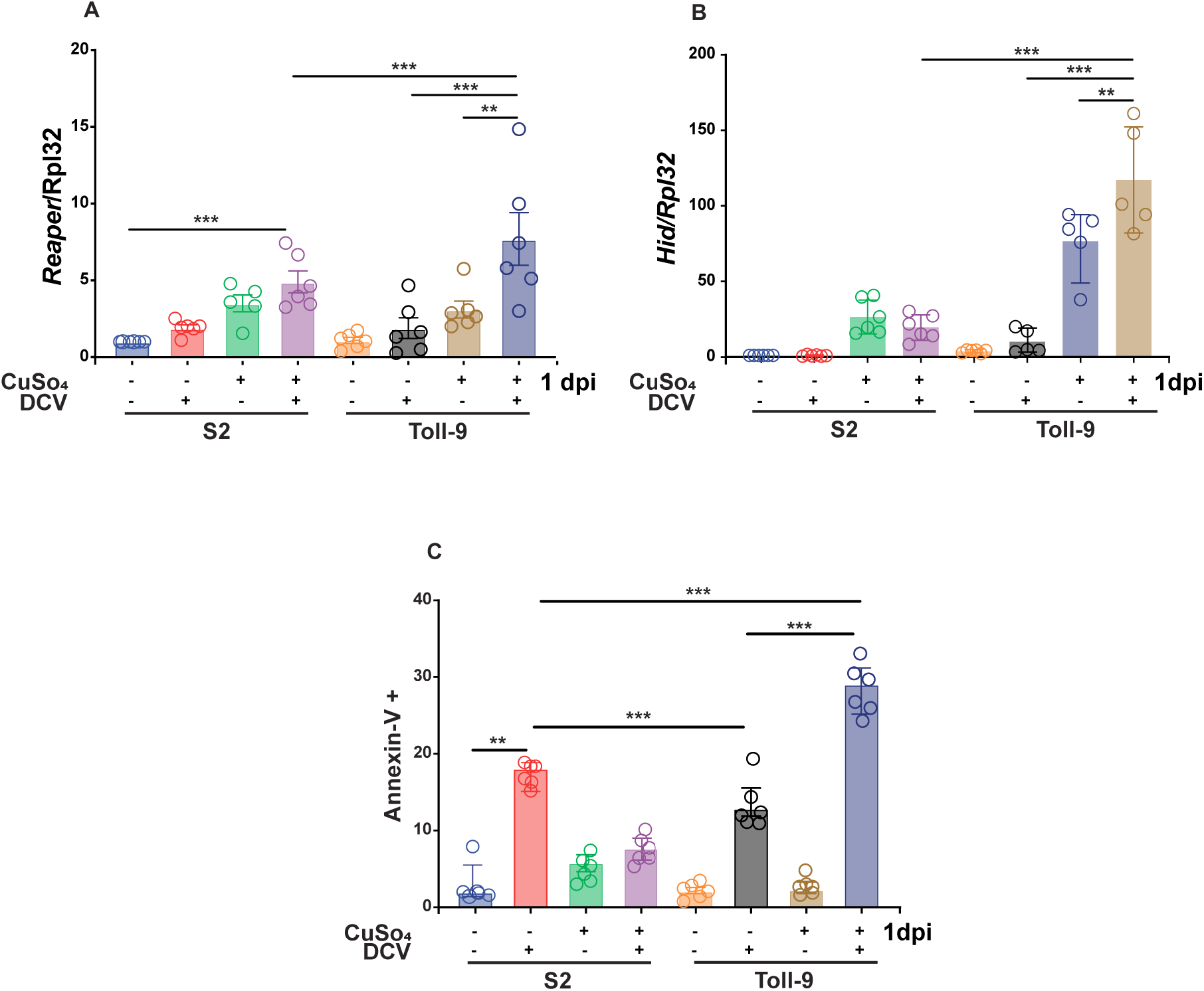
Toll-9 induces apoptosis during DCV infection of S2 cells. **(A-B)** qPCR analysis DCV infected S2 naive and S2 cells expressing TOll-9 in presence and absence of CuSO_4_ at 1 dpi to detect proapoptotic genes *Reaper* and *Hid*. **(C)** Percentage of Annexin V positive cells from DCV infected S2 naive and S2 cells expressing TOll-9 in presence and absence of CuSO_4_ at 1 dpi. Data are representative of six biological replicates (well of cells) from three independent experiments. Error bars, SEM. Unpaired T test, **p<0.01; ***p<0.001.

## DISCUSSION

The structural similarity observed between Toll-9 and mammalian Toll-like receptors (TLRs), alongside their recognized involvement in innate immune mechanisms, prompted the hypothesis that Toll-9 may play a role in innate immunity in *Drosophila* (Ooi et al. 2002). However, Narbonne-Reveau et al. showed that the loss of Toll-9 does not affect basal antimicrobial peptide (AMP) levels or their induction during bacterial infection (Narbonne-Reveau, Charroux, and Royet 2011). Nevertheless, recent studies highlight significant roles for Toll-9 in cell competition, aging, and neurodegeneration. Specifically, Toll-9 mediates apoptosis in unhealthy cells and promotes proliferation in fit cells (Alpar, Bergantiños, and Johnston 2018; Shields et al. 2022; Sakakibara et al. 2023). Despite these advances, Toll-9’s role in viral infections, its specific activator ligand, and the mechanistic details of its signaling cascade remain elusive and warrants further investigation.

DCV is one of the most studied natural pathogen of *Drosophila*, having coevolved with its host (Jousset et al. 1972; Sabin, Hanna, and Cherry 2010). During infection, DCV activates various host immune pathways, including phagocytosis, RNAi, and JAK/STAT signaling (Sabin, Hanna, and Cherry 2010; Karlikow, Goic, and Saleh 2014; Nainu et al. 2015). Given the structural similarities between Toll-9 and human TLRs, we hypothesized that it is involved in the antiviral response against DCV. Our study demonstrates Toll-9’s role in the antiviral response to the DCV infection in adult flies and S2 cells. A recent study using *Bombyx mori* Toll-9 (BmToll-9) reveals that it shares structural similarities with mammalian TLR4 and functions as pattern recognition receptor (PRR) that detects PAMP lipopolysaccharide (LPS) (R. Zhang et al. 2021). Previous research on *B*. *mori* immune related genes has identified two Toll-9 variants with immune functions. Among these, BmToll-9-1 is more closely related to *Drosophila* Toll-9 and plays crucial role in gut immune response (Wu et al. 2010). BmToll-9-1 expressing Bm5 cells, when exposed to dsRNA, exhibit significant upregulation of the key RNAi component gene *Dicer2*, highlighting a pivotal role for BmToll-9-1 in activating the RNAi pathway (Liu et al. 2014). RNAi is a crucial antiviral mechanism in arthropods (Mussabekova, Daeffler, and Imler 2017), and it consists of two primary arms mediated by Dicer1 and Dicer2. Dicer1 plays a role in both RNA degradation and translational repression processes, whereas Dicer2 is primarily involved in RNA degradation. The loss of RNAi components *Dicer2, Argonaute2,* or *r2d2* increases susceptibility to viral infection (Galiana-Arnoux et al. 2006). Our investigation revealed that DCV-infected transposable element insertion mutant *Toll-9^0024-G4^* flies show reduced induction of RNAi components such as *Dicer2* and *Argonaute2,* leads to the higher mortality rates compared to infected isogenic parental control *w^1118^* flies. Contrary to expectations, our investigation revealed higher induction levels of JAK/STAT pathway components, including *Upd2, Upd3*, and *Vir1*, in mutant *Toll-9^0024-G4^*flies compared to isogenic parental control *w^1118^* flies. This finding suggests that while the JAK/STAT pathway was upregulated, it alone was insufficient to provide adequate protection to the flies in the absence of induction of the RNAi pathway, as previously observed (Dostert et al. 2005).

Many viruses promote host cell proliferation to prolong their replication and evade immune clearance, often through phosphorylation of AKT (Datta, Brunet, and Greenberg 1999; Ji and Liu, n.d.; Wei et al. 2012; Cooray 2004). For example, in *Drosophila* S2 cells, Sindbis virus enhances AKT phosphorylation, which supports virus replication (Patel and Hardy 2012). Insulin-mediated PI3K/AKT signaling also regulates the expression of RNAi genes *Dicer2* and *Argonaute2* via FoxO during West Nile and Zika virus infection (Ahlers et al. 2019; Trammell et al. 2022). Our study reveals that Toll-9 expression during DCV infection of S2 cells restricts AKT phosphorylation, thereby reducing viral load. Notably, to our knowledge, this is the first study to examine AKT’s role in DCV replication in *Drosophila* S2 cells. Toll-9 inhibits AKT activation through its dephosphorylation, resulting in the upregulation of *Dicer2* and reduction of DCV replication in Toll-9 expressing S2 cells. A recent study on *Drosophila* dopaminergic neuronal cell survival shows that overexpression of Toll-1 and Toll-7 induces autophagy by dephosphorylating AKT via activation of PP2A (Protein Phosphatase 2A) suggesting for the role of Toll signaling in regulating phosphorylation of AKT (J. Zhang et al. 2024). Furthermore, pharmacological inhibition of AKT during DCV infection in S2 cells mimics the restrictive effect of Toll-9, underscoring Toll-9’s role in inhibiting DCV infection through AKT signaling.

Mammalian TLRs exhibit diverse subcellular localization, with TLRs 1, 2, 4, 5, and 6 being primarily localized on the cell membrane interacting with extracellular PAMPs, while TLRs 3, 7, 8, 9, and 10 are predominantly found in endosomes or lysosomes where they sense intracellular PAMPs such as nucleic acids (Gay et al. 2014; S. M.-Y. Lee et al. 2018: El-Zayat, Sibaii, and Mannaa 2019). Our findings indicate that *Drosophila* Toll-9 localizes to both early and late endosomes, suggesting a role in nucleic acid sensing. This hypothesis is further supported by Toll-9’s ability to bind to poly(I:C), a double-stranded RNA analogue, implying that it can detect viral dsRNA, similar to human TLR10 (S. M.-Y. Lee et al. 2018). During DCV infection, the detection of dsRNA by the Toll-9 in endosomes triggers signaling that inhibits AKT phosphorylation. This signaling leads to upregulation of proapoptotic genes and reduction of viral loads within the cell through apoptosis (Jeong et al. 2008). This parallels the mechanism in mammalian system where TLRs mediate immune responses, underscoring the evolutionary significance of Toll-9 in antiviral defenses.

Apoptosis is a form of programmed cell death essential for development, homeostasis and defense against pathogens (Kerr, Wyllie, and Currie 1972). Studies indicate that inhibition of AKT phosphorylation in HTLV-1 transformed cells induces apoptosis by decreasing phosphorylated Bad, leading to cytochrome C release and caspase-9 activation. (Jeong et al. 2008). Similarly, in streptozotocin-induced diabetic mice, reduced phosphorylation of AKT induces apoptosis via cytochrome C release and increased caspase-3 activity (Meng et al. 2017). In *Drosophila*, DCV infection triggers apoptosis in S2 cells by reduction of *Drosophila* inhibitor of apoptosis (DIAP1) that inhibits caspases (Nainu et al. 2015; Nainu, Shiratsuchi, and Nakanishi 2017). Recent research demonstrates that Toll-9, in conjunction with Toll-1, activates the intracellular Toll-1 pathway to induce apoptosis and promote the expression of proapoptotic genes such as *Hid* and *Reaper* during cellular surveillance, facilitating the elimination of compromised cells (Meyer et al. 2014; Alpar, Bergantiños, and Johnston 2018, Shields et al. 2022). During *Drosophila* development, *Hid* and *Reaper* activate Dronc (a caspase-9 homolog), which then activates DrICE and Dcp-1 (caspase-3 and caspase-7 homologs), leading to cell death (Shalini et al. 2015; Salvesen, Hempel, and Coll 2016). However, the mechanism through which apoptosis is triggered in S2 cells during viral infection is not fully understood. Here, we show that in DCV infected Toll-9 expressing S2 cells, Toll-9 induces AKT dephosphorylation, which upregulates *Hid* and *Reaper* that could be antagonizing DIAP1, leading to Dronc and DrICE activation and apoptosis initiation. Overactivation of the PI3K/AKT pathway is a hallmark of many human cancers where it inhibits apoptosis and promotes cell survival (Fu and Tindall 2008; X. Zhang et al. 2011). Studies show that pharmacological inhibition of AKT inhibition induces apoptosis via nuclear retention of FoxO (Roy, Srivastava, and Shankar 2010; Das et al. 2016; Sun et al. 2018; N. Lee et al. 2020). Our findings suggest Toll-9 mediates apoptosis in DCV infected S2 cells by inducing the expression of DIAP1-antagonizing genes, *Hid* and *Reaper*, likely through the AKT dephosphorylation.

In conclusion, we demonstrated that Toll-9 is crucial for antiviral defense against DCV infection in S2 cells. Toll-9 restricts DCV replication by inducing expression of the RNAi pathway gene *Dicer2* and restriction of AKT phosphorylation leading to apoptosis. Our data demonstrate that Toll-9 is localized in endosome similar to human TLR10 and can bind to dsRNA. This investigation will shed light on Toll-9’s potential role in modulating cellular immune responses against viral infections, providing valuable insights into its function in the innate immune system of *Drosophila*.

## MATERIALS AND METHODS

### Fly stocks and insect cell culture

*Drosophila melanogaster* genotypes used were *w^1118^*and *w^1118^*; *PBac{w^+mC^=IT.GAL4} Toll-9^0024-G4^* (Bloomington *Drosophila* Stock Center 5905 and 62581). Flies were grown in standard meal agar fly food and maintained at 23°C and 68% humidity under a 12h/12h light/dark cycle. *Drosophila* hemocyte-derived S2 cells were maintained at 28°C in tissue culture flasks containing Schneider’s *Drosophila* medium (Gibco, Waltham, MA) supplemented with 10% heat-inactivated fetal bovine serum (FBS) (HyClone), 100 U/ml of penicillin, 100 μg/ml of streptomycin, and 0.25 μg/ml of amphotericin B (Fungizone) antimycotic (Life Technologies, Waltham, MA).

### DCV infection

Age matched four-seven days old flies were infected with DCV (5000 TCID_50_) or mock (Saline), and mortality was evaluated for a period of 30 days. For injections, flies were anesthetized with CO_2_ and injected with 23 nl of virus or PBS using a pulled 0.53-mm glass needle and an automatic nanoliter injector (Drummond Scientific, Broomall, PA). Individual flies were injected at the ventrolateral surface of the fly thorax and placed into new vials. After the injections, the adult flies were monitored daily for mortality and collected at different times post-infection to assess viral load or host responses. Survival curves were performed using a minimum of 80 flies per condition, including at least three experimental replicates. The viral titer was analyzed observing the cytopathic effect of serially diluted cell culture supernatant from DCV infected S2 cells by calculating TCID_50_/ml.

### Gene expression analysis

The relative expression of each RNAi and JAK/STAT pathway genes was determined in adult flies by quantitative reverse transcription PCR (qRT-PCR). Six biological replicates of S2 cells or five biological replicates of five flies each per condition were lysed in Trizol Reagent (ThermoFisher 15596), and RNA was isolated by column purification (ZymoResearch R2050) and cDNA was synthesized using the iScript Reverse Transcriptase kit (Bio-Rad, Hercules, CA). qPCR was performed using SYBR green PCR master mix (Bio-Rad).

*RpL32* (5′-GACGCTTCAAGGGACAGTATCTG-3′ and 5′-AAACGCGGTTCTGCATGAG-3′) *Hid* (5-ACGGCCATCCGAATCCGAAC-3 and 5-TGCTGCTGCCGGAAGAAGAAGTT-3), *Reaper* (5-GATGGTCTGGATTCCATTGC-3 and 5-TAGTCTGCGCCAACATCATC-3) *Vir1* (5’-GATCCCAATTTTCCCATCAA-3’ and 5’-GATTACAGCTGGGTGCACAA-3’), DCV (5’-TCATCGGTATGCACATTGCT-3’ and 5’-CGCATAACCATGCTCTTCTG-3’), *Ago2* (5’-CCGGAAGTGACTGTGACAGATCG-3’ and 5’ CCTCCACGCACTGCATTGCTCG-3’) *Upd2* (5’-CCTATCCGAACAGCAATGGT-3’ and 5’ CTGGCGTGTGAAAGTTGAGA-3’), *Upd3* (5’-GCCCTCTTCACCAAACTGAA -3’ and 5’ TCGCCTTGCACAGACTCTTA-3’), *Dcr2* (5’-GTATGGCGATAGTGTGACTGCGAC -3’ and 5’ GCAGCTTGTTCCGCAGCAATATAGC-3’),

### Plasmid constructs and transfection into S2 cells

The coding regions of *Drosophila* Toll-9 (CG5528) were amplified from *Drosophila* Genomics Resource Center clone IP19811. Amplified PCR fragments were inserted into the metallothionein promoter driven pMT-V5-His vector (Invitrogen). The generated vector was co-transfected with a pCoBlast blasticidin resistance vector with a ratio of 19:1 and cells were selected in blasticidin containing medium according to manufacturer’s instructions. After establishing several stably transfected lines, the expression level was checked with CuSO_4_ induction (500 μM) and the cells were used for the experiments performed in this study.

### Immunoblotting

Flies or S2 cells were homogenized in RIPA buffer (25 mM Tris-HCl [pH 7.6], 150 mM NaCl, 1 mM EDTA, 1% NP-40, 1% sodium deoxycholate, 0.1% SDS, 1 mM Na3VO4, 1 mM NaF, 0.1 mM phenylmethylsulfonyl fluoride [PMSF], 10 μM aprotinin, 5 μg/ml leupeptin, 1 μg/ml pepstatin A). Total protein was determined using the bicinchoninic acid (BCA) assay (Pierce, Waltham, MA). Equal amounts of protein were subjected to SDS-PAGE. The proteins were transferred to a polyvinylidene difluoride (PVDF) membrane and blocked in 5% bovine serum albumin (BSA) in 0.1% Tween 20–Tris-buffered saline. The membrane was incubated with antibodies against DCV capsid (1:1000; Abcam ab92954), V5 tag (1:5000; ThermoFisher Scientific R960-25), actin (1:5000; Sigma A2066), and rab5 (1:1000; Abcam ab31261) antibodies overnight at 4°C. Antibody-bound proteins were detected using anti-rabbit secondary antibodies conjugated to horseradish peroxidase. The blots were developed by chemiluminescence using luminol enhancer solution (ThermoFisher).

### Fluorescence microscopy

The infected cells were fixed for 20 min in 4% formaldehyde, followed by permeabilization for 20 min in 0.1% Triton X-100. The cells were blocked in phosphate-buffered saline (PBS) containing 10% FBS and incubated with antibodies against Rab5 (1:50; Abcam ab31261), Rab7(1:20; Developmental Studies Hybridoma Bank Rab7), DCV Capsid (1:200; Abcam ab92954), dsRNA J2 (1:200; Jena Biosciences RNT-SCI-10010200) and V5 tag (1:200; ThermoFisher Scientific R960-25) for overnight at 4°C. The cells were washed and incubated with rabbit or mice Alexa Fluor-488 (ThermoFisher Scientific A11034 or A11035)- and rabbit or mice Alexa Fluor-594 (ThermoFisher Scientific A11029 or A11030)-conjugated secondary antibodies for 1 h at room temperature. The cells were washed, incubated with DAPI (4′,6-diamidino-2-phenylindole) (Sigma D9542) for 15 min, and mounted onto microscope slides. Images were obtained using a Leica DMi8 fluorescent microscope and analyzed using Leica Application Suite X.

### Poly (I:C) delivery and immunoprecipitation

To deliver poly (I:C) to the S2 cells or Toll-9 expressing S2 cells, 10^6^ cells were incubated with 5 μg/ml poly (I:C) as described previously (Li and Zamore 2019). Cell culture media was removed before the treatment of poly (I:C) and replaced with the incomplete cell culture media without FBS to starve the cells for 1 hour to increase delivery of poly (I:C). Incomplete cell culture media was replaced with complete cell culture media and cells were allowed to grow for 48 hours followed by induction with 500 μM CuSO_4_ for 24 hours. The cells were harvested and lysed using RIPA buffer with protease inhibitors. For poly (I:C), pulldown 200 μg lysate and incubate with 2 μg anti-dsRNA J2 antibody for 16 hours at 4°C on rotating nutator. After incubating the cell lysate and anti-dsRNA J2 antibody, 40 μl of 50% v/v protein A agarose beads were added to the mixture and incubated for 3 hours at 4°C on rotating nutator. Bead-antibody-protein complexes were washed 3 times in 1X RIPA buffer and then incubated for 10 minutes at 97 °C with 2X loading dye. Lysates were centrifuged for 5 minutes at 13,000 × g to remove the beads and subjected for western blotting analyses.

### Flow cytometry

Annexin V binding buffer, fluorochrome-conjugated Annexin V, and propidium iodide (Invitrogen A35110) were used according to manufacturer’s instructions. The cells were detached using gentle pipetting and were washed once with PBS and once with Annexin V binding buffer. Cells were stained with Annexin V (1:200 dilution) for 15 min at room temperature in Annexin V binding buffer. Data was acquired using Easycyte Guava flow cytometer and analyzed using Guava soft software.

### Organelle isolation

To isolate cytosol and microsome, cells were washed twice with PBS at 300g for 5 minutes at 4℃. After washing once with PBS, cells were washed with 1 mL 0.9% sodium chloride solution at 300 x g for 5 minutes at 4℃. The pellet was resuspended in the lysis buffer provided in the kit and cells were fractionated using Qproteome-mitochondria isolation kit according to manufacturer’s instructions (Qiagen 37612).

### Quantification and statistical analysis

An unpaired two-tailed Student’s t test assuming unequal variance was utilized to compare means of quantitative data. Mortality curves were analyzed by the log-rank (Mantel-Cox) test using GraphPad Prism (GraphPad Software, Inc.). Results shown are representative of at least three independent experiments. Circles in dot plots represent individual biological replicates of a well of cells of groups of five flies.

## Acknowledgments

We are thankful to Ziying Liu, and Miyoung Lee for experimental support for fly experiments.

This research was supported by NIH/National Institute of Allergy and Infectious Diseases (NIAID) (grant R01 AI139051 to A.G.G.).

Conceptualization by M.C. and A.G.G.; methodology by M.C. designed and performed the experiments and wrote the paper; B.M.H and P.E.M. performed the fly injections and qRT PCR experiments and revised the paper; M.C and A.G.G. conceived the study, designed experiments, supervised the experiments, and revised the paper.

